# The Cyclin-Like Protein Spy1 Mediates Tumourigenic Potential of Triple Negative Breast Cancer

**DOI:** 10.1101/2024.03.11.584461

**Authors:** Bre-Anne Fifield, Claudia Pecoraro, Amy Basilious, Catalin Gramisteanu, Emily Mailloux, Rosa-Maria Ferraiuolo, Lisa A. Porter

## Abstract

Triple negative breast cancer is an aggressive subtype of breast cancer that relies on systemic chemotherapy as its primary means of treatment. Cell cycle regulators are enriched in drug resistant forms of the disease supporting the potential of targeting cell cycle checkpoints as a therapeutic direction to re-sensitize patients to treatment. Spy1 is an atypical cyclin-like protein that can override cell cycle checkpoints and is elevated in triple negative breast cancer. We report for the first time the effects of CRISPR-Cas9 mediated knockout of Spy1 on functional characteristics of triple negative breast cancer cells and perform unbiased analysis of protein expression to assess global changes in expression which correlate with functional changes in cell properties. Loss of Spy1 reduced rates of proliferation, decreased metastatic potential, and led to a reduction in stemness properties of triple negative breast cancer cells. Importantly, knockout of Spy1 delayed tumour onset in an *in vivo* model and significantly increased response to chemotherapy, pushing cells towards a senescent state. This data reveals that changes in expression of proteins that are not essential for proliferation and only transiently expressed can have significant impacts on cell dynamics and provides support for targeting the Spy1-CDK2 complex as a new therapeutic avenue in triple negative breast cancer.

**Statement of Significance:** Targeting the atypical cell cycle regulator Spy1 induces senescence and increases responsiveness of triple negative breast cancer to standard of care chemotherapy.

## Introduction

Triple negative breast cancer (TNBC) accounts for approximately 15 to 20% of all breast cancer diagnoses (1). This form of cancer progresses independently of estrogen, progesterone and Her2 signaling, rendering it non-responsive to targeted therapies that have greatly benefited patients with other subtypes of breast cancer. TNBC therefore relies on systemic chemotherapy, alongside radiation and surgery. Following treatment, approximately 40% of TNBC patients will have a pathological complete response which correlates with improved disease free and overall survival (2). The remaining 60% of patients with residual disease are at increased risk of disease progression and relapse (3). Understanding the molecular regulators mediating this response is of utmost importance to improve treatment outcomes for this high-risk group of patients.

Data supports that residual disease is associated with alterations in expression of cell cycle regulators and DNA repair pathways and has an increased proportion of breast cancer stem cells (BCSCs), a drug resistant population of cells capable of self-renewal (4–6). Increased expression of select cyclins has been associated with residual disease and supports propagation of BCSC populations (5,6). Elevated cyclin expression corresponds to increased activity of their binding partners, cyclin dependent kinases (CDKs), which would allow for enhanced cell cycle progression. In combination with decreased expression of cyclin dependent kinase inhibitors (CKIs) such as p16, p27 and p21 which work to inhibit cyclin-CDK complexes, this would render cells insensitive to chemotherapy which often relies on activation of cell cycle checkpoints to push cells towards apoptosis (7,8). Recent work has explored cell cycle vulnerabilities as a means of targeted therapy by examining essentiality of cell cycle regulators across cell lines (9). Dependency on G1/S regulators CDK4, CDK6, CDK2 and cyclins E1 and D1 differ widely between specific cell lines studied (9). Cell lines dependent on cyclin E1 are more sensitive to depletion of epigenetic and DNA replication factors, revealing that understanding these cell cycle vulnerabilities may lead to targeted therapy approaches (9,10). The focus, however, has been on regulators deemed essential for proliferation, or genes which are significantly amplified in TNBC and does not consider transient increases in expression of mediators that can override checkpoints. While not essential for proliferation, these genes can have significant effects on cell cycle dynamics. Transient increases in expression may be one mechanism by which cells are able to evade chemotherapy and represent an important area of study. A full understanding of the mechanisms regulating these checkpoints may reveal novel ways to reinstate them and represent a powerful therapeutic option to sensitize tumours to therapy.

One often overlooked family of cell cycle regulators is the Speedy/RINGO family (originally identified member Spy1; gene SPDYA). Spy1 can activate CDK2 independent of post-translational modifications required for activation of classic cyclin-CDK complexes, rendering it insensitive to inhibition by CKIs (11–16). Through this unique mechanism of CDK2 activation, Spy1 can drive cell cycle progression and override a variety of checkpoints (12,17–19). Unlike canonical cyclins, Spy1 levels are low in most somatic cells, except at specific periods of development (12,20,21). Previous work has demonstrated that levels of Spy1 are increased in TNBC, exogenous elevation of Spy1 can increase resistance to various forms of treatment and transient knockdown supports the potential to sensitize different cancers to therapy (22–24). Thus, targeting of Spy1 may reinstate checkpoints and increase sensitivity to therapy, improving treatment responses for patients who may otherwise not respond.

Herein, we explore the effects of knockout of Spy1 for the first time in a cell line model of TNBC and perform unbiased analysis to obtain a snapshot of the overall changes to protein expression that may result from loss of this unique cell cycle regulator. Our results have revealed global changes in expression of regulators of proliferation and apoptosis as well as overall decreases in expression of poor prognostic indicators in TNBC. Loss of Spy1 alters functional properties of TNBC cells, including decreased proliferation and a decreased proportion of BCSCs. This data also reveals that loss of Spy1 delays tumour onset in an *in vivo* model and significantly increases response to therapy. Together, this preclinical data provides exciting support for the potential of select targeting of Spy1 as a therapeutic avenue in TNBC.

## Materials and Methods

### Cell Line Data

DepMap data was downloaded from the DepMap data portal (<u>DepMap Data Downloads</u>). The Project Score, CERES dataset containing information for 318 cancer cell lines was utilized and CERES scores were calculated as described (25,26).

### Cell Culture

MDA-MB-231 (HTB-26; ATCC) were cultured in Dulbecco’s modified Eagle’s medium (DMEM; D5796; Sigma Aldrich) supplemented with 10% foetal bovine serum (FBS; 35-077-CV; Corning) and 1% P/S at 37°C and 5% CO_2_. A BioRad TC10 Automated Cell Counter was used to assess cell viability via trypan blue exclusion. MDA-MB-231 cells were lentivirally infected in serum and antibiotic free medium containing 8µg/mL polybrene. DHB-mVenus (CDK2 activity reporter) was a gift from Tobias Meyer & Sabrina Spencer (Addgene plasmid # 136461; http://n2t.net/addgene:136461; RRID:Addgene_136461) (27).

### Drug Treatment

Dimethyl sulfoxide (DMSO) was used as the vehicle control for all treatments. Cells were treated with 25nM doxorubicin (Sigma), 6mM cyclophosphamide (Sigma), 100nM paclitaxel (Thermo), and 50µM carboplatin (Sigma). All dosages used were IC50s based off trypan blue exclusion analysis and MTT assay as described (22).

### Generation of Spy1 Deletion Using CRISPR

Guide RNAs were designed using the CRISPR Design Tool (http://tools.genome-engineering.org). The CRISPR-Cas9 cloning vector px459 (Addgene 62988; (28)) was a kind gift from Fred Dick, Lawson Health Research Institute. Forward and reverse primers (100 µM) were phosphorylated and annealed using T4 Polynucleotide Kinase (PNK; New England Biolabs M0201S) and 10x T4 Ligation Buffer (Thermo B69). The px459 vector (25ng) and phosphorylated and annealed oligos were digested and ligated in one reaction using FastDigest BbsI (Thermo FD1014), T4 DNA ligase (Thermo 15224017), and T4 ligase buffer (Thermo B69). 30,000 cells were seeded in a 24-well plate 24 hours prior to transfection. Cells were transfected with 1 µg of branched PEI and 400 ng of DNA. PEI and DNA were incubated in 100 µL of growth media with no Pen/Strep or serum for 10 minutes before being added to the cells. The day following transfection, the media was removed and replaced with full growth media. Cells were given 24 hours to recover before the media was removed and replaced with growth media containing 1 µg/mL puromycin to select for transfected cells. Cells were allowed to recover before being single cell sorted into a 96 well plate for clonal selection using the BD FACSAria Fusion™ 5. After clonal selection, successful deletion of Spy1 was confirmed via sequencing and western blot analysis.

### Flow Cytometry

MDA-MB-231 control and Spy1 knockout cells were stained with propidium iodide after collection during an exponential growth phase. DNA content was assessed using the BD LSR Fortessa™ X-20.

### Mammosphere Formation Assay

MDA-MB-231 cells were seeded at a density of 2500 cells per mL of media (DMEM/F-12 medium (Corning 10-092-CV), 1% Pen/Strep, 20 ng/mL basic fibroblast growth factor (bFGF; Sigma F0291), 20 ng/mL EGF, and 100 µg/mL gentamicin (Thermo 15710064)). Primary spheres were measured and quantified 7 days after seeding.

### Reverse Phase Protein Array (RPPA)

MDA-MB-231 control or Spy1 knockout cells were subjected to RPPA analysis as previously described (29,30). A total of 488 antibodies were included in the analysis. Relative protein levels for each sample were determined by RPPA space (31). Presented heat maps were generated using Morpheus (https://software.broadinstitute.org/morpheus/). Proliferation and apoptotic scores were calculated from mean expression level (log2 scale) taking the average expression of positive regulators and subtracting the average expression of negative regulators of each process (32).

### Immunoblotting

Cell pellets were lysed as previously described (17). Bradford assay was performed as per manufacturer instructions to assess protein concentrations and equal amounts of protein were analysed and separated using SDS-PAGE and transferred to PVDF membranes. Membranes were blocked at room temperature for 1 hour and incubated in primary antibody overnight at 4°C. Chemiluminescent Peroxidase Substrate was used for visualization following manufacturer’s instruction (Pierce) and quantified on an AlphaInnotech HD2 (Fisher) using AlphaEase FC software. Primary antibodies were used as follows: tubulin (1:1000; Cell Signaling 2146S); actin (1:1000; Millipore MAB1501R), Spy1 (1:1000; Thermo PA5-29417), integrin β1 (1:1000; Cell Signaling 4706); integrin α5 (1:1000; Cell Signaling 4705); cyclin D3 (1:1000; Cell Signaling 2936S), cyclin B1 (1:2000; Thermo MA1-155), cyclin D1 (1:1000; Abcam ab16663); Sox2 (1:1000; Abcam ab97959).

### Quantitative Real-Time PCR Analysis

RNA was isolated from cell pellets using Qiagen RNeasy Kit and cDNA was synthesized using Quanta qScript following manufacturer’s instructions. SYBR Green detection (Applied Biosystems) was used for real-time PCR. Viia7 Real Time PCR System (Life Technologies) and software was used for PCR and analysis.

### Adhesion Assay

24-well plates were pre-coated with collagen type I (5µg/mL, Corning 354236) or fibronectin (10µg/mL, Sigma F2006) for 1 hour at room temperature. MDA-MB-231 control and Spy1 knockout cells were seeded on untreated plates and plates pre-coated with collagen type I and fibronectin at a cell density of 20,000 cells per well. Plates were placed in an incubator for 1hr, 4hr and 8hr timepoints. After incubation, non-adherent cells were washed with PBS. Adherent cells were fixed using ice cold 100% methanol and stained using a 0.5% crystal violet stain. Images were taken on a microscope (Leica DM IL LED) and analyzed using ImageJ analysis.

### Invasion and Migration Assay

Transwell migration and invasion assays were performed using 12-well 8µm cell inserts which were placed into 12-well plates with 1mL of full growth media (DMEM, 10% FBS, 1% P/S). The upper chambers were untreated, precoated with Cultrex (R&D Systems 3433-005-01), or collagen type I (5µg/mL, Corning 354236) and placed in an incubator for 30 minutes. MDA-MB-231 control or Spy1 knockout cells were seeded at a cell density of 100,000 cells per well in 500µL of serum free media (DMEM) and placed in the incubator for 24 hours. After incubation, the inserts were washed in PBS, cells were fixed using 4% PFA and stained using a 0.5% crystal violet stain. Images were taken on a microscope (Leica DM IL LED) and analyzed using ImageJ analysis.

### Xenograft Transplants

MDA-MB-231 control or Spy1 knockout cells were orthotopically injected into the inguinal mammary glands of 8-week-old NOD/SCID (Jackson lab; 001303). 500,000 cells per gland were injected and allowed to grow until tumour endpoint was reached. Mice were maintained following the Canadian Council on Animal Care Guidelines under animal utilization protocol 20-15 approved by the University of Windsor.

### Immunohistochemistry

Tumour tissue was fixed in 10% neutral buffered formalin. Immunohistochemistry was performed as described (33). Primary antibodies were diluted in 3% BSA-0.1% Tween-20 in 1x PBS. Primary antibodies were used as follows: cleaved caspase 3 (1:250; Cell Signaling) and phospho-histone H3 (1:300; Abcam) both for 1hr at room temperature. Secondary biotinylated anti-rabbit (Vector Laboratories) was used at a concentration of 1:750. Slides were imaged use a Leica DMI6000 inverted microscope and quantification performed using ImageJ.

### Senescence Associated Beta-Galactosidase Staining

Cells were fixed in 4% PFA followed by incubation overnight at 37°C in pre-warmed X-gal at a final concentration of 1mg/mL in staining solution (5mM potassium ferrocyanide, 5mM potassium ferricyanide, 2mM MgCl2; pH6). The following day, staining solution was removed, cells were washed with 1x PBS and visualized under a Leia DMI6000 inverted microscope.

### Statistical Analysis

A Mann-Whitney test was performed for statistical analysis on tumour studies. For all other data, a Student’s T Test was performed. Analysis for cell line data assumed equal variance, and mouse tissue sample data assumed unequal variance. All experiments include 3 biological and technical replicates.

## Results

### Loss of Spy1 delays cell cycle progression and reduces CDK2 activity

The dependency of breast cancer cell lines on Spy1 was first explored using DepMap data (25,26) in which CRISPR-Cas9 deletion was used to test essentiality of genes as indicated by the CERES score. A CERES score less than −1 typically indicates a given gene is considered essential for proliferation (9,25,26). While the average score never reached −1, TNBC cell lines had the lowest CERES score when compared to hormone receptor or Her2 positive cells (Fig 1A). When examining a small panel of commonly used TNBC cell lines, the MDA-MB-231 cells had the lowest CERES score (Fig 1B). To explore the impact of loss of Spy1 on cell dynamics of MDA-MB-231 cells, CRISPR-Cas9 was used to induce Spy1 knockout. Successful knockout was confirmed via western blot analysis (Fig 1C) as well as sequencing analysis. Knockout of Spy1 lead to a significant decrease in proliferative capacity with a reduction in total number of live cells as assessed via a trypan blue exclusion assay (Fig 1D), as well as a significant increase in doubling time (Fig 1E). Cell cycle profile was assessed in control versus Spy1 knockout cells via flow cytometry. Loss of Spy1 led to a significant increase in the percentage of cells in G1 and S phase and a reduction in cells in G2/M phase (Fig 1F). Since Spy1-mediated proliferation is controlled, at least in part, via direct interaction and activation of CDK2 (13), Spy1 knockout cells were assessed using the DHB-mVenus CDK2 activity reporter (27). For this system, localization of the mVenus reporter is used to assess CDK2 activity, with nuclear localization indicating low CDK2 activity (27). The percentage of cells with nuclear versus cytoplasmic localization of the DHB-mVenus reporter was quantified. Spy1 knockout cells had a significant increase in cells with low CDK2 activity and a significant decrease in cells with high CDK2 activity (Fig 1G). Thus, knockout of Spy1 decreases the proliferative capacity of MDA-MB-231 cells in part by decreasing CDK2 activity, demonstrating that deletion of genes not evaluated as essential may still have significant impacts on cell dynamics.

**Figure 1:**
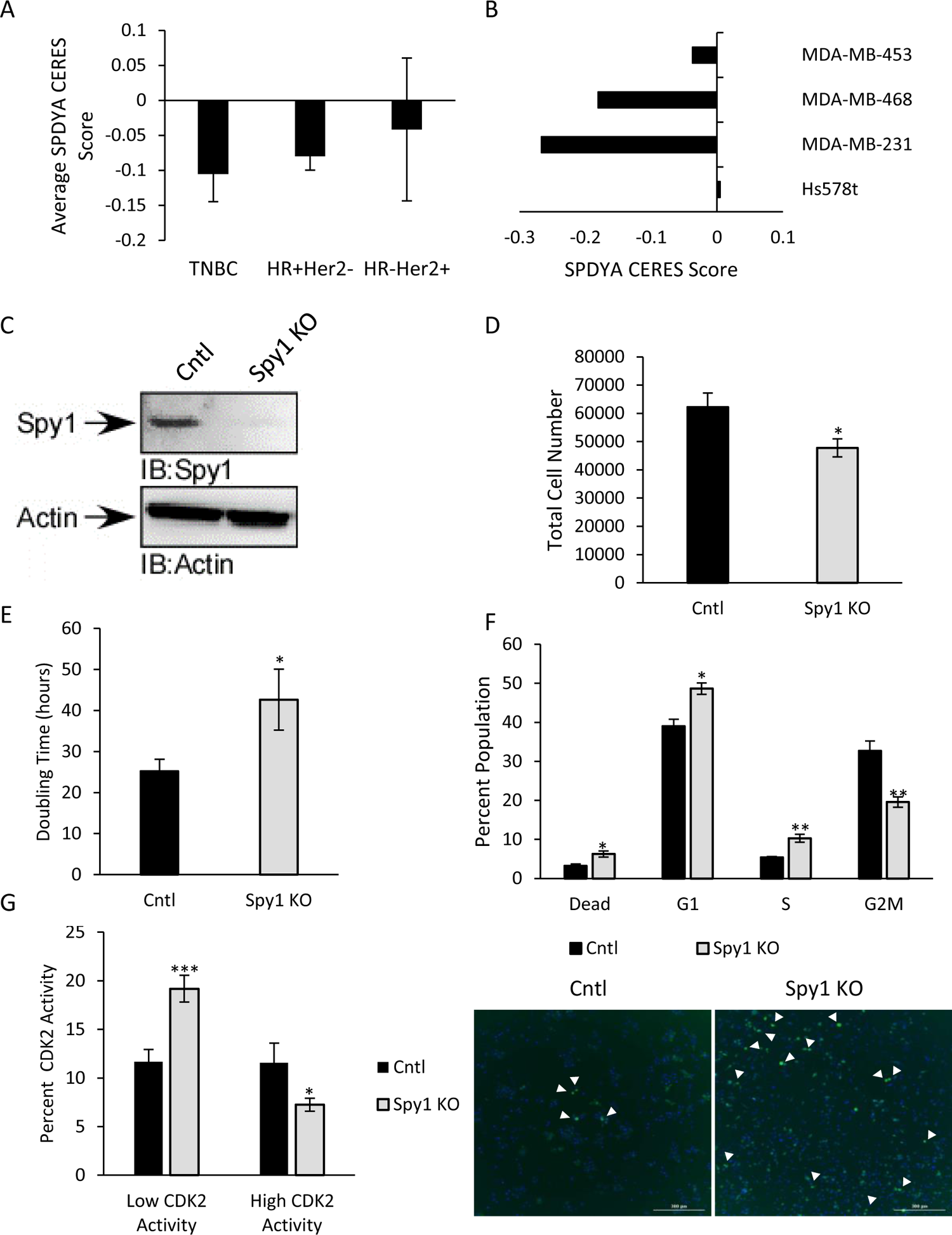
Loss of Spy1 decreases proliferative potential. DepMap was used to assess CERES scores across **A)** cell lines of varying breast cancer subtypes and **B)** common triple negative breast cancer cell lines. **C)** Levels of Spy1 protein were assessed via western blot analysis. Representative blot is depicted. **D)** Growth of MDA-MB-231 control versus Spy1 knockout cells was assessed via trypan blue exclusion assay. Total number of live cells is depicted graphically. **E)** Doubling time in hours of MDA-MB-231 control and Spy1 knockout cells. **F)** Cell cycle distribution of MDA-MB-231 control and Spy1 knockout cells was assessed via flow cytometry. Percentage of cells in each phase of the cell cycle is depicted. **G)** MDA-MB-231 control and Spy1 knockout cells were infected with the DHB-mVenus CDK2 activity reporter. Representative images in right panel and left panel depicts percentage of cells with low or high CDK2 activity. White arrowheads point to nuclear localization of signal. n=3; Error bars represent SE; Student’s T test. *p < 0.05, **p < 0.01, ***p < 0.001

### Spy1 loss reduces expression of proteins involved in cell proliferation and stemness and increases apoptotic proteins

To determine if other cyclin and Speedy/RINGO family members could be compensating for the loss of Spy1, qRT-PCR analysis was performed. There were no significant differences in the mRNA expression of Speedy/RINGO family members or cyclins examined (Fig S1), thus indicating any changes observed are likely due to the loss of Spy1 protein. Protein lysates from MDA-MB-231 control and Spy1 knockout cells were sent for RPPA analysis for an unbiased screen of changes to protein expression upon loss of Spy1. Cell cycle, DNA damage and senescence proteins were assessed for changes in protein expression (Fig 2A). Loss of Spy1 resulted in changes in protein expression across the spectrum of proteins assessed. Western blot analysis was used to verify a select set of cell cycle proteins to validate the RPPA data, and validated the trends observed with RPPA (Fig 2B). Proliferation and apoptotic scores were calculated from RPPA data considering both positive and negative regulators of proliferation and apoptosis (Fig S2, S3). MDA-MB-231 Spy1 knockout cells had a decreased proliferation score (Fig 2C) and an increased apoptotic score (Fig 2D) indicative of decreased proliferation and increased apoptosis as was seen in functional assays (Fig 1).

**Figure 2:**
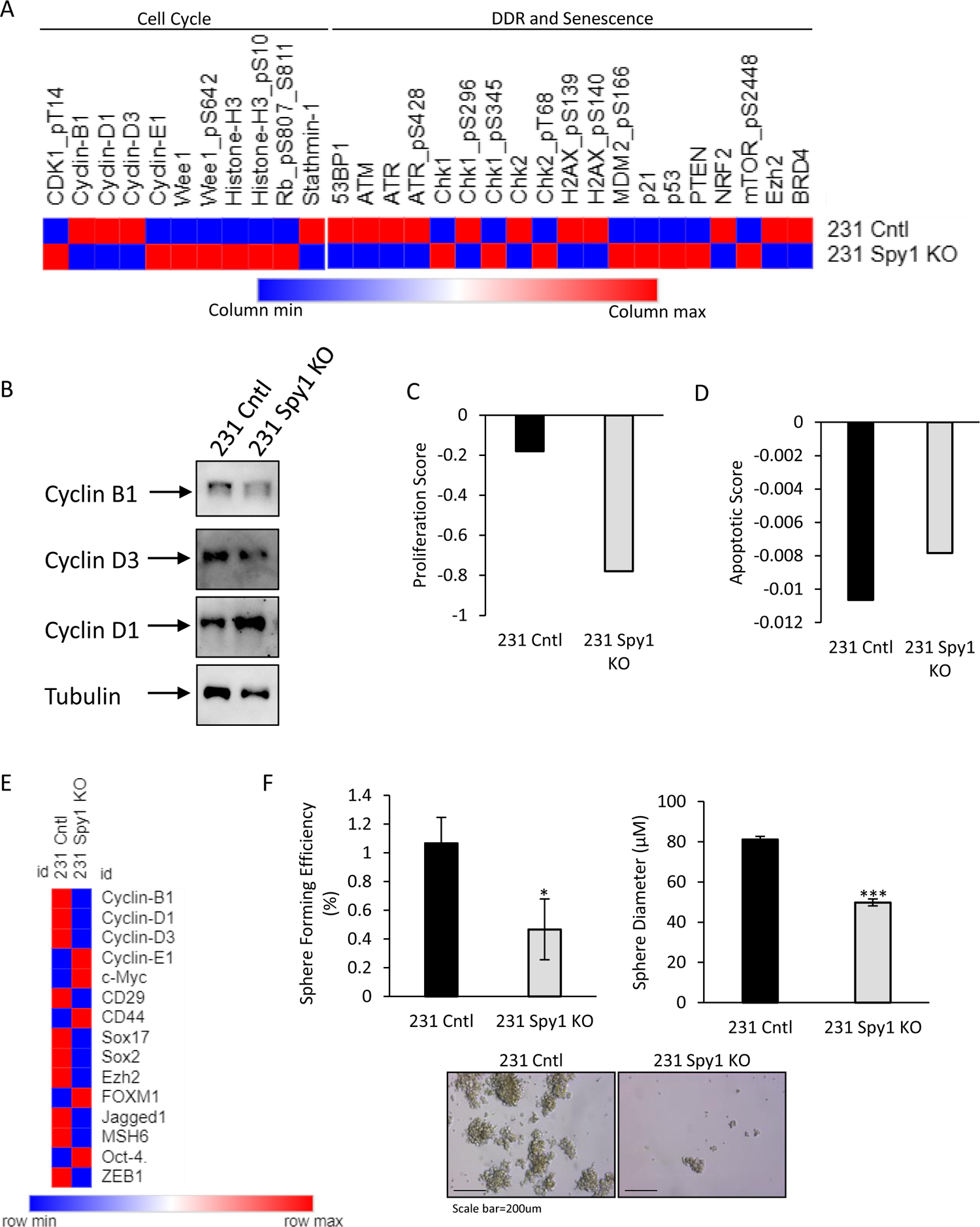
Alterations in protein expression profiles with loss of Spy1 correlate with functional assays. **A)** Heat map of log2 expression profiles of select cell cycle, DNA damage response (DDR) and senescent markers from RPPA expression analysis of MDA-MB-231 control and Spy1 knockout cells. **B)** Levels of select cell cycle mediators were assessed via western blot analysis to validate RPPA findings. Representative blot is shown (n=3). **C)** Proliferation and **D)** apoptotic score of MDA-MB-231 control and Spy1 knockout cells calculated from log2 mean expression of cell cycle and apoptotic regulator expression from RPPA analysis. **E)** Log2 expression profile of markers correlating with breast cancer stem cells based on RPPA analysis. **F)** MDA-MB-231 control and Spy1 knockout cells were seeded for a mammosphere formation assay. Sphere forming efficiency (top left panel) and size of mammospheres (top right panel) are depicted graphically. Representative images are shown in bottom panel (n=3). Error bars represent SE; Student’s T test. *p < 0.05, **p < 0.01, ***p < 0.001

### Spy1 loss reduces stemness properties of TNBC

Based on this data, select mediators of BCSC population expansion were assessed (Fig 2E). While there were inconsistencies in changes in expression of proteins which are known to expand the BCSC population, overall, the trend demonstrated a decrease in expression of positive regulators of the BCSC population (Fig 2E). To functionally verify this finding, MDA-MB-231 cells were seeded in sphere formation assays. Knockout of Spy1 resulted in a significant decrease in sphere forming efficiency with significantly smaller spheres (Fig 2F). This data demonstrates that loss of Spy1 results in changes in protein expression which have functional effects on not only proliferation but also stemness potential.

### Spy1 contributes to metastatic potential of TNBC

Using the RPPA data, an expression profile of mediators of metastatic potential in TNBC was generated (Fig 3A). In general, most of these proteins were downregulated upon Spy1 knockout (Fig 3A). To functionally validate these findings, the adhesion properties of MDA-MB-231 cells with Spy1 knockout were assessed on tissue culture plates (Fig S4), as well as with collagen type 1 and fibronectin (Fig 3B). Loss of Spy1 significantly decreased adhesion to all substrates assessed (Fig 3B). Next, migration and invasion of Spy1 knockout cells was analysed. While loss of Spy1 did not significantly decrease migration (Fig 4A), it did significantly decrease invasion through Cultrex (Fig 4B). Expression of key mediators of the cellular interaction between collagen type I and IV (integrin β1) and fibronectin (integrin α5) were examined via western blot analysis (Fig 4C). The knockout of Spy1 resulted in decreases in integrin β1 and integrin α5 (Fig 4C). Thus, Spy1 may play a role in metastasis of TNBC.

**Figure 3:**
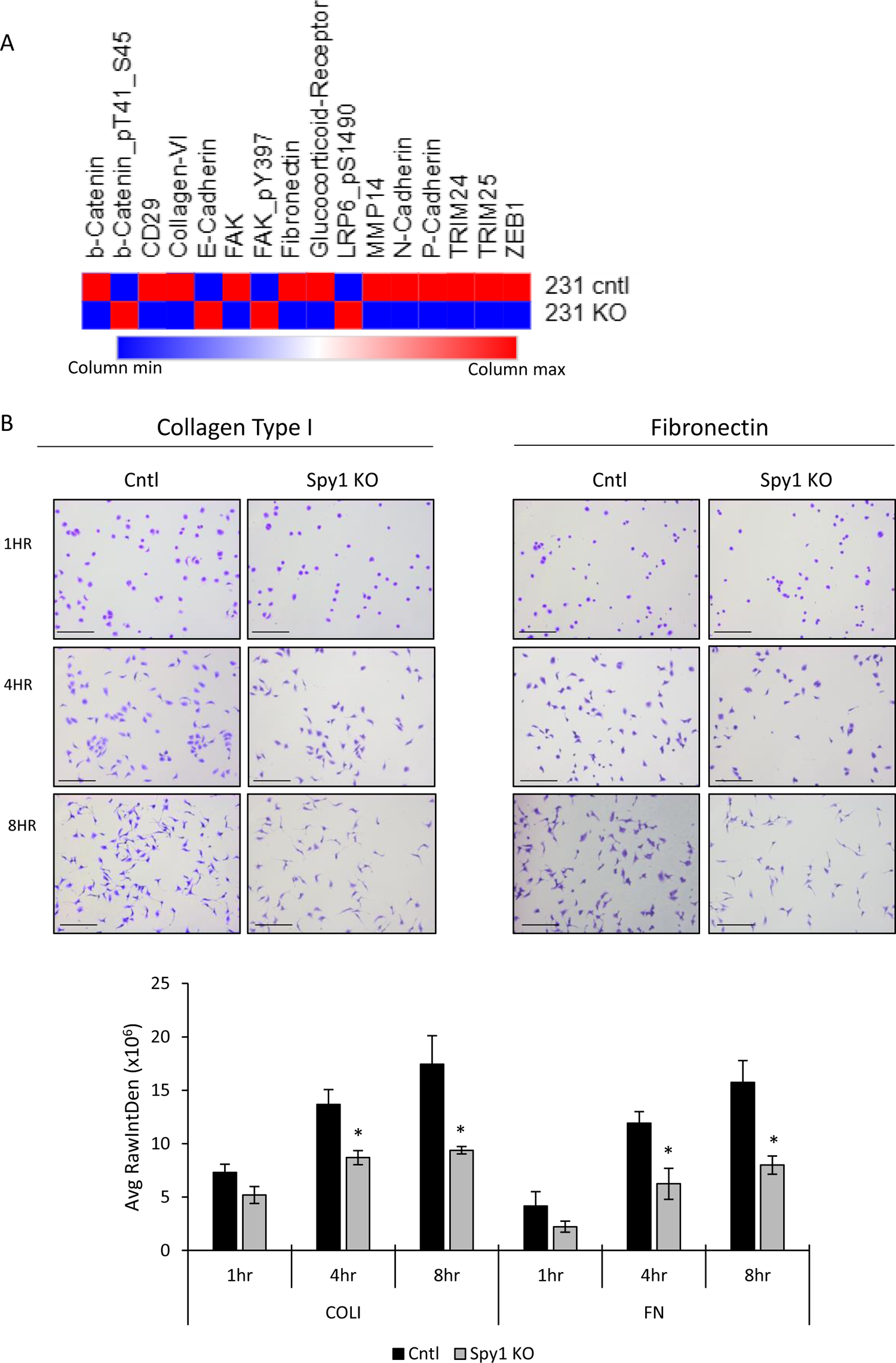
Spy1 mediates adhesion of TNBC cells. **A)** Heat map depicting log2 expression of mediators of metastasis from RPPA analysis. **B)** Representative images (top panel) of MDA-MB-231 adhesion assay of Spy1 knockout and control cells on either collagen type I (COLI) or fibronectin (FN) coated plates at 1hr, 4hr and 8hr timepoints. Images were taken at 10X magnification. Scale bar represents 200µm. Graph (bottom panel) represents quantification of MDA-MB-231 Spy1 knockout cells compared to MDA-MB-231 control cells adhered to the plate at respective timepoints. n=3. Error bars represent SE; Student’s T test. *p < 0.05, **p < 0.01, ***p < 0.001

**Figure 4:**
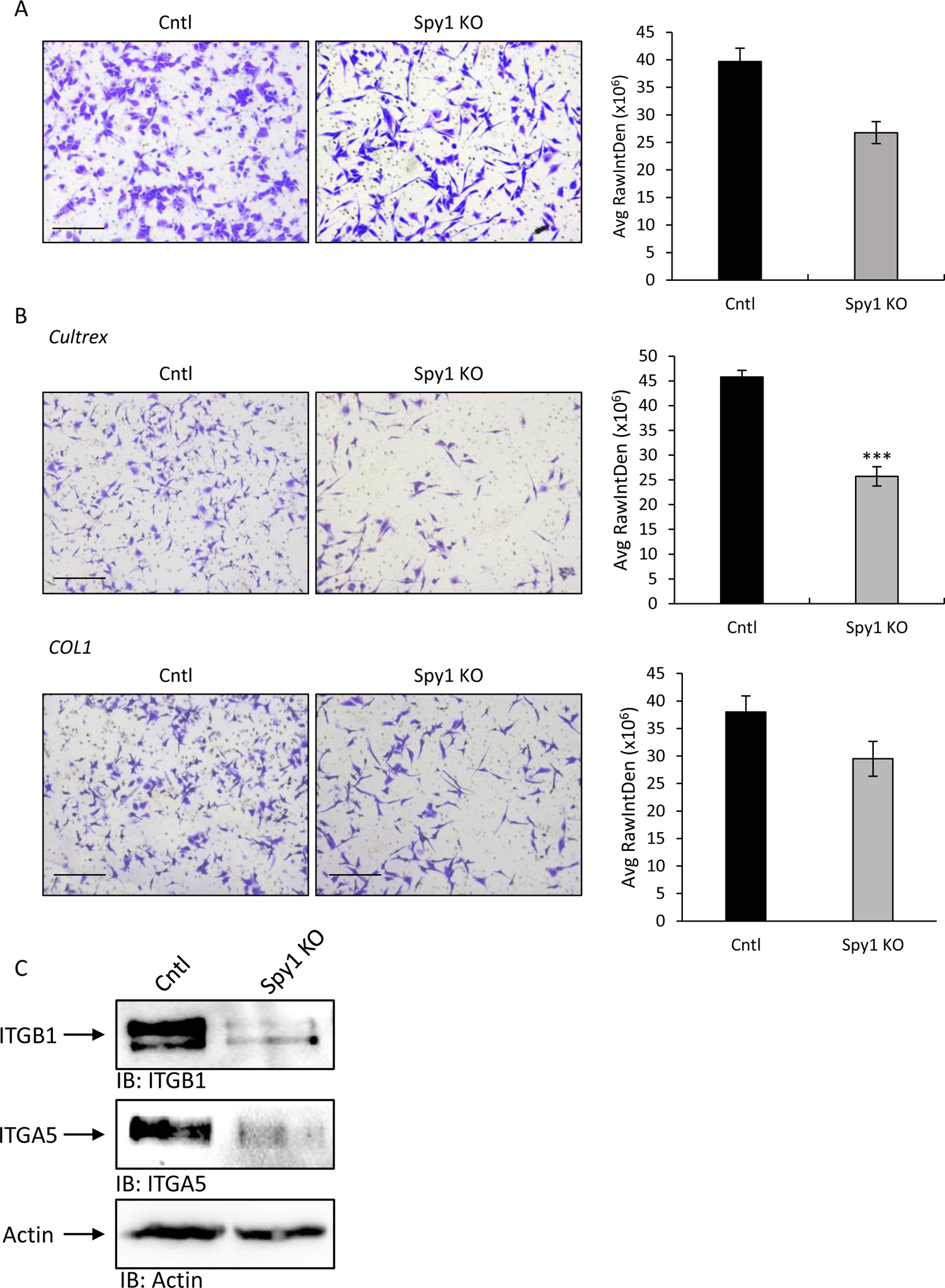
Decreased invasive properties and altered integrin expression with loss of Spy1. **A)** Migration assay, and invasion assays on **B)** Cultrex or **C)** COL1 were conducted on MDA-MB-231 control or Spy1 knockout cells. Representative images are depicted in left panels and quantification is depicted graphically in right panels. Images were taken at 10X magnification. Scale bar represents 200µm. **C)** Levels of select integrin proteins were assessed via western blot analysis. Representative blot is shown. n=3. Error bars represent SE; Student’s T test. *p < 0.05, **p < 0.01, ***p < 0.001

### Knockout of Spy1 delays tumour onset and increases sensitivity to therapy

Decreased proliferative capacity and metastatic potential could indicate improved prognosis upon loss of Spy1 expression in TNBC. Expression of indicators of TNBC prognosis were examined using the RPPA dataset and demonstrated that many of these indicators were decreased with Spy1 knockout (Fig 5A). To determine if this translates to decreased tumourigenic potential, MDA-MB-231 control and Spy1 knockout cells were orthotopically transplanted into 8-week-old NOD/SCID mice and monitored weekly for tumour development. Even with a small sample size, knockout of Spy1 delayed tumour development as compared to control cells, and tumours from Spy1 knockout cells were on average smaller than control tumours (Fig 5B, C). Immunohistochemical analysis revealed a significant increase in apoptosis in Spy1 knockout tumours as assessed by the percentage of positive cleaved caspase 3 cells (Fig 5D). Interestingly, despite a smaller overall tumour size and increased apoptosis, Spy1 knockout tumours also had a significant increase in phospho-histone H3 positive cells (Fig 5E), a trend that was also observed in the RPPA data set from MDA-MB-231 Spy1 knockout cells (Fig 2A). Since Spy1 knockout tumours retained the same trend with phospho-histone H3, western blot analysis was then performed to determine if changes in protein expression observed in the cells were conserved in tumour tissue. Expression of integrin β1, integrin α5, cyclin D3 and Sox2 were all decreased in Spy1 knockout tumours as compared to control (Fig S5), mimicking what was observed in the MDA-MB-231 Spy1 knockout cells.

**Figure 5:**
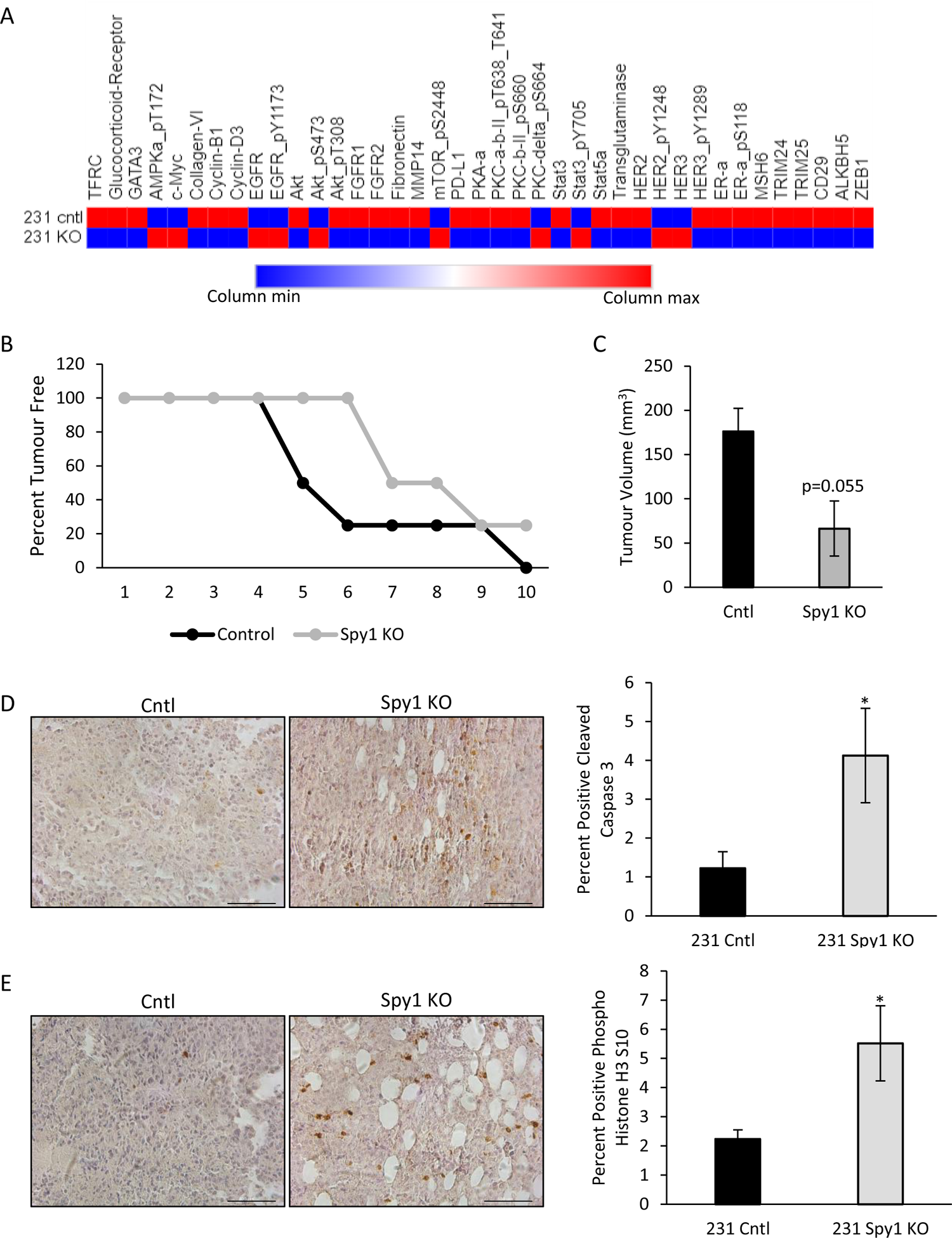
Knockout of Spy1 delays tumourigenesis. **A)** Heat map depicting log2 expression profile of prognostic markers for TNBC from RPPA analysis of MDA-MB-231 control and Spy1 knockout cells. **B)** MDA-MB-231 control and Spy1 knockout cells were orthotopically injected into inguinal mammary glands of 8-week-old NOD/SCID mice. Mice were monitored on a weekly basis for tumour formation (n=4). Graphical representation of percent of tumour free mice. **C)** Tumours were measured using calipers once per week. Tumour volume of control or Spy1 knockout tumours is depicted graphically. Immunohistochemical analysis of **D)** cleaved caspase 3 and **E)** phospho-histone H3 in control and Spy1 knockout tumours where blue stain represents hematoxylin and brown stain represents **D)** cleaved caspase 3 or **E)** phospho-histone H3 (left panels). Percent positive **D)** cleaved caspase 3 and **E)** phospho-histone H3 cells is depicted graphically in right panels (n=3). Scale bar=100µM. Error bars represent SE; Student’s T test. *p < 0.05

To further characterize the impact of loss of Spy1 on tumourigenic potential, MDA-MB-231 control and Spy1 knockout cells were treated with standard of care chemotherapy used for TNBC, either doxorubicin and cyclophosphamide or paclitaxel and carboplatin, and total live cells quantified 24 and 48 hours post treatment. Loss of Spy1 significantly decreased the total number of live cells at both time points assessed for both treatments (Fig 6A). The induction of senescence was also quantified 1 week after 24 hours of treatment with standard of care (Fig 6B). Spy1 knockout cells had a significant increase in senescence following treatment with the combination of doxorubicin and cyclophosphamide (Fig 6C). Expression of CKIs involved in cell cycle arrest and induction of senescence were then examined via qRT-PCR. Expression of p21 was significantly increased in both control and Spy1 knockout cells 48hrs post-treatment, but then significantly decreased in Spy1 knockout cells by 1 week post treatment (Fig S6A); however, a significant increase in expression of p16 was observed in Spy1 knockout cells 1 week following treatment, and no change in p16 expression was seen in MDA-MB-231 control cells (Fig 6D). Overall, this supports that loss of Spy1 not only sensitizes cells to therapy but could also be used for induction of senescence for novel therapeutic approaches (Fig 7).

**Figure 6:**
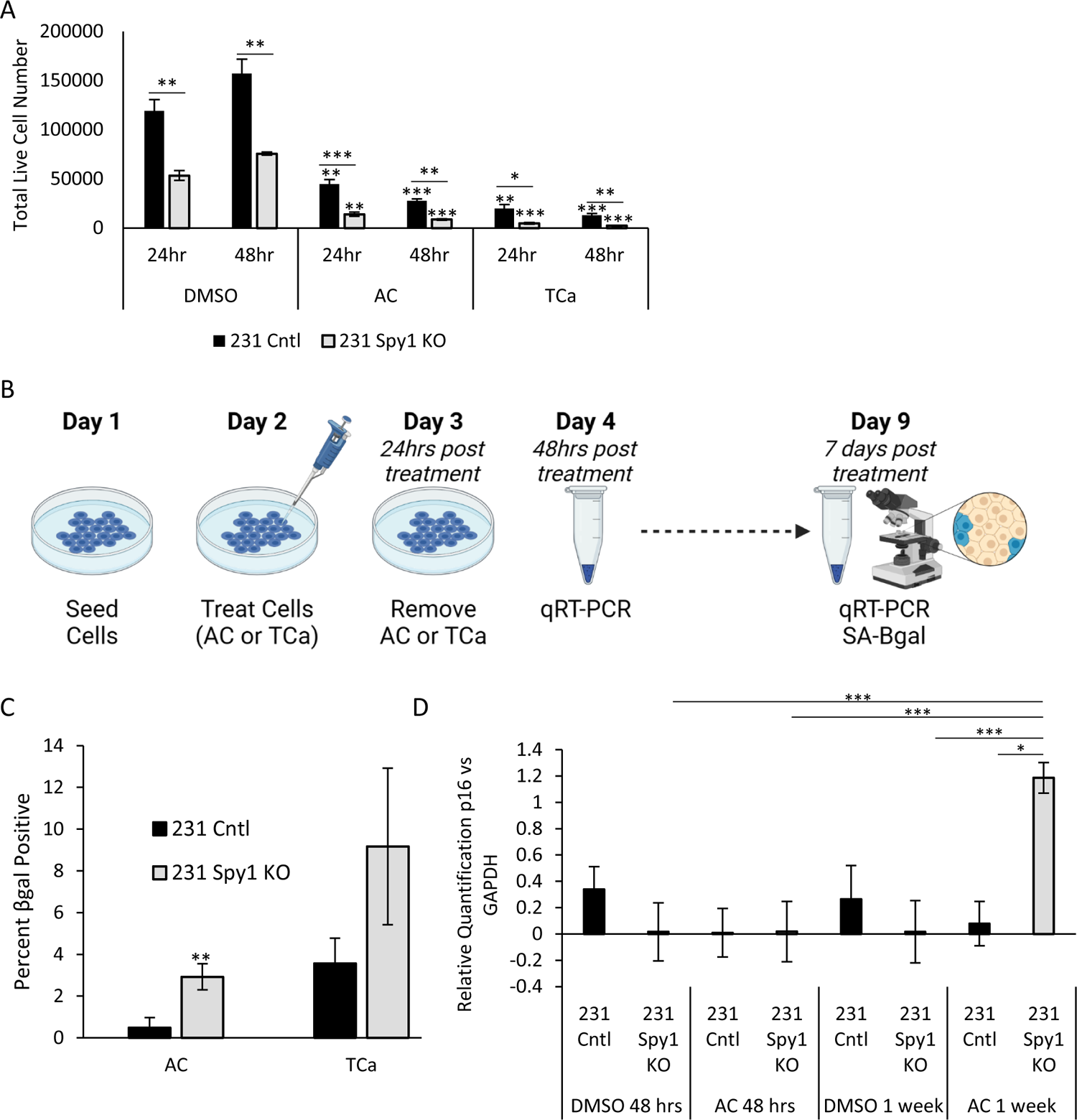
Spy1 contributes to therapy resistance. **A)** MDA-MB-231 control and Spy1 knockout cells were treated with doxorubicin and cyclophosphamide (AC) or paclitaxel and carboplatin (TCa) for 24 and 48 hours. Total live cells were quantified via trypan blue exclusion analysis. **B)** Treatment schema of MDA-MB-231 control and Spy1 knockout cells treated with AC or TCa and followed for CKI expression and induction of senescence. **C)** Quantification of senescence associate β-galactosidase (βgal) expression in MDA-MB-231 control and Spy1 knockout cells 1 week following treatment with AC or TCa. Percentage of cells positive for βgal staining is depicted graphically. **D)** qRT-PCR analysis of p16 levels corrected for GAPDH on MDA-MB-231 control and Spy1 knockout cells treated with AC. n=3. A=doxorubicin, C=cyclophosphamide, T=paclitaxel, Ca=carboplatin; Error bars represent SE; Student’s T test. *p < 0.05, **p < 0.01, ***p < 0.001

**Figure 7.**
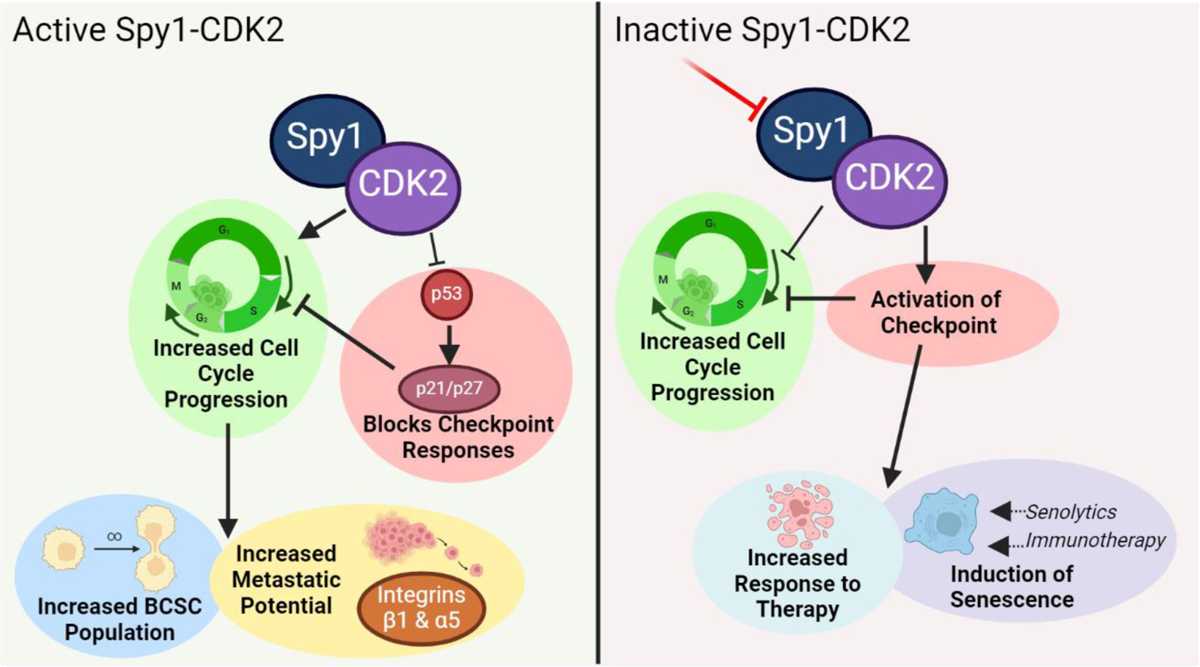
Schematic of functional changes observed with loss of Spy1 in triple negative breast cancer.

## Discussion

Enrichment of cell cycle regulators has been found in patients who are non-responsive to chemotherapy, thus finding novel ways of re-instating cell cycle checkpoints could prove a valuable therapeutic option for many patients at high risk of relapse. While much work has focused on canonical cell cycle regulators that are essential drivers of proliferation as a means of cell cycle targeting, this work demonstrates that even regulators which may be transiently expressed may prove to be a valuable therapeutic approach. Knockout of Spy1 significantly reduced rates of proliferation in the TNBC cell line MDA-MB-231 and decreased overall CDK2 activity within the heterogenous population. When orthotopically transplanted into NOD/SCID mice, loss of Spy1 delayed tumour onset and resulted in overall smaller tumours with increased apoptosis. These changes were seen despite the fact Spy1 was not considered an essential gene for proliferation based on the CERES score obtained from the DepMap portal. This may be due to redundancies in CDK2 activation as in the context of loss of Spy1, cyclins A or E may compensate and allowed for continued proliferation at reduced rates.

An unbiased screen of alterations in protein expression in Spy1 knockout cells was conducted to determine if there were any large-scale changes in expression that are associated with loss of Spy1 that could reveal mechanistic insight into the role of Spy1 in TNBC. The average expression of cell cycle and apoptotic regulators were used to generate a proliferation and apoptotic score which matched the functional changes observed in cell growth and cell cycle assays indicating these types of scores may have the potential to assess proliferative potential when assessing alterations or use of new therapeutics. Expression of mediators of different pathways associated with prognostic potential of TNBC were also examined. Decreased expression of proteins associated with metastatic potential was observed with loss of Spy1. This corresponded with a significant reduction in adhesion, migration and invasion, all important processes in the metastatic cascade. Additionally, a reduction in integrin β1 and integrin α5 expression was also observed. Both integrin β1 and integrin α5 have been associated with poor prognosis in TNBC and reduction of their expression leads to reduced metastatic potential (34,35). Further work examining the role of Spy1 in metastasis is of high priority to determine the potential of targeting Spy1 to reduce metastatic burden in TNBC patients.

Interestingly, integrin β1 has also been linked to the BCSC population, where loss of integrin β1 leads to a reduction in self-renewal capacity of this population (36). Knockout of Spy1 also lead to a reduction in the BCSC population and reduction in expression of other mediators implicated as drivers of this dangerous population of cells. Previous work has demonstrated that Spy1 promotes a switch to symmetric division in the CD133 positive population of brain tumour initiating cells (37). Whether Spy1 has the same role in BCSCs or is working through another mechanism is an important area of future study. A decrease in the BCSC population may also lead to an increase in sensitivity to therapy, as this population is a highly drug resistant population of cells enriched in residual disease (4). When treated with standard of care chemotherapy for TNBC, Spy1 knockout cells had a significant reduction in overall total live cell number as compared to control cells indicating increased sensitivity to therapy. Previous work has demonstrated that Spy1 can override a variety of cell cycle checkpoints, including those activated in response to DNA damage, much like would be induced by various chemotherapeutic agents (11,17–19,22). This provides evidence that targeting Spy1 could help to re-instate cell cycle checkpoints.

Cyclin E1 amplification has been associated with poor prognosis in TNBC (38) and depleting cyclin E1 reduces tumourigenic potential in both TNBC cell line models and mouse models of breast cancer (39). Tumours with amplified cyclin E1 have also been reported to be vulnerable to WEE1 inhibitors, targeting the high levels of replication stress seen with cyclin E1 overexpression (9,10). Understanding vulnerabilities associated with either amplification or loss of expression could reveal new combination therapies for more targeted approaches. In addition to increased sensitivity to chemotherapy, knockout of Spy1 also significantly increased senescence following treatment. Inducing senescence as a means of therapy has recently gained attention as a valuable therapeutic approach (40). Triggering senescence may help to sensitize aggressive cell populations to immunotherapy approaches or novel strategies to directly target senescent cells, referred to as senolytics (41–43). This is of particular importance as treatment of TNBC now includes the use of the immunotherapy agent pembrolizumab; however, even with this approach, approximately 20% of patients will relapse within 3 to 5 years (44,45).

This work is the first to explore the effects of loss of Spy1 on functional properties of TNBC. Loss of Spy1 reduced CDK2 activity and overall rates of proliferation and significantly reduced the dangerous population of BCSCs. Knockout of Spy1 reduced the overall metastatic potential and delayed tumour onset resulting in smaller tumours with increased apoptosis. Importantly, loss of Spy1 re-sensitized cells to treatment and increased senescence, providing evidence to support targeting of Spy1 as a therapeutic option. The unique structure and mechanism of the Spy1-CDK2 complex could prove to be a druggable target that could help to re-instate checkpoints, increasing sensitivity to already existing treatment protocols. Additionally, this mechanism could be targeted as means to improve response to immunotherapy approaches and new approaches targeting senescent populations. Given its low levels of expression in somatic tissues, this targeted approach may also lead to less side effects than targeting canonical cyclin-CDK complexes which are widely expressed. Thus, novel approaches targeting the Spy1-CDK2 mechanism could help improve outcomes and open new avenues of treatment options for those with aggressive, treatment resistant forms of TNBC.

## Acknowledgements

Special thanks to the University of Windsor Central Animal Care Facility and University of Windsor Flow Cytometry Facility, and Dr. F. Dick for the px459 vector. Thanks to Drs E. Fidalgo da Silva and D. Lubanska, as well as I. Hinch and N. Philbin for technical assistance. The Functional Proteomics Reverse Phase Protein Array Core was supported in part by The University of Texas MD Anderson Cancer Center, P30CA016672, and R50CA221675. This work was supported by operating funds from the Canadian Institutes Health Research to L.A.P (Grant#142189).

## Data Availability

The data generated in this study are available upon request from the corresponding author.

