## Supplemental Files for "The Cyclin-Like Protein Spy1 Mediates Tumourigenic Potential of Triple Negative Breast Cancer"

Figure S1

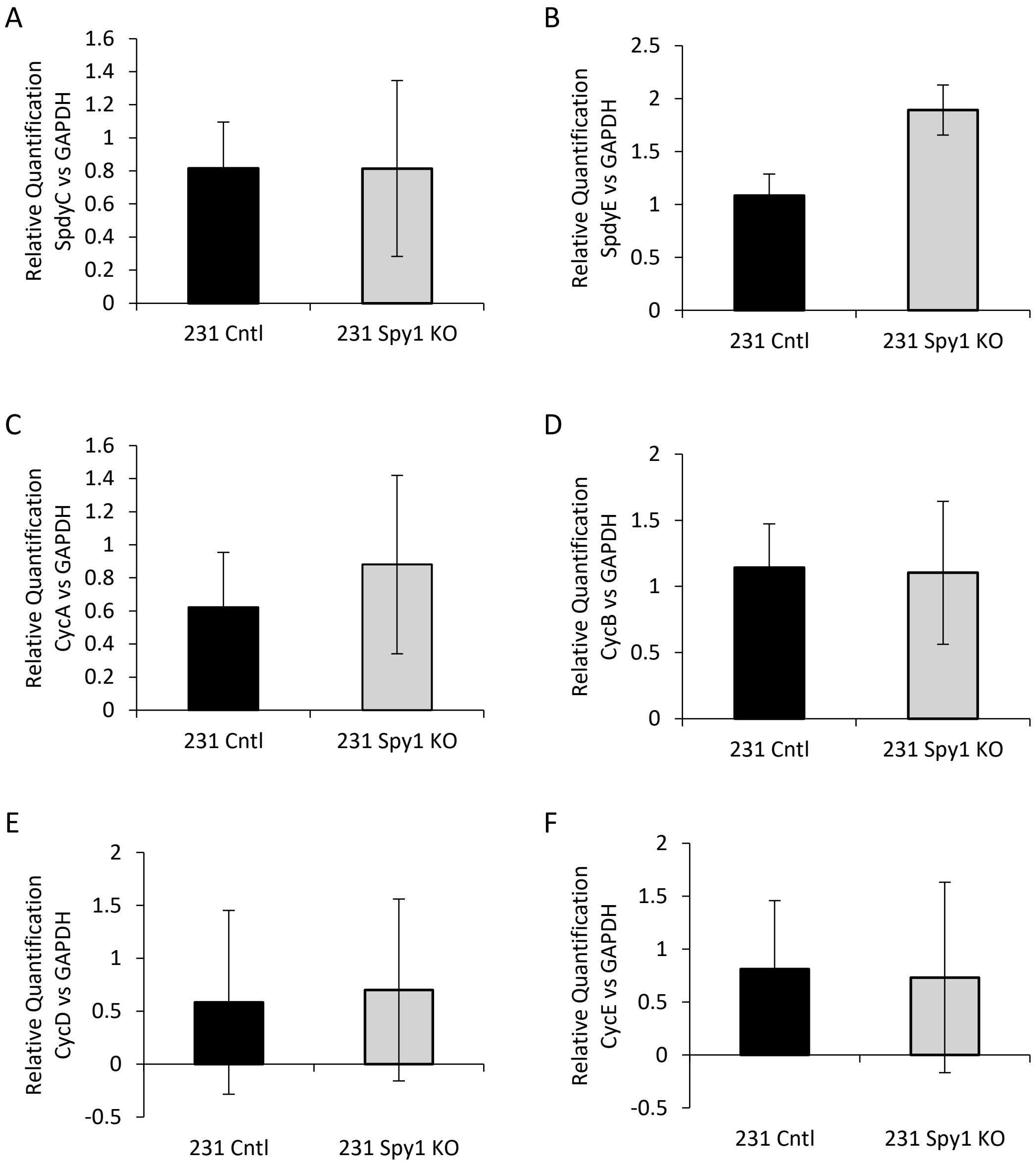
A
Relative Quantification SpdyC vs GAPDH
1.6
1.4
1.2
1
0.8
0.6
0.4
0.2
0
231 Cntl
231 Spy1 KO
B
Relative Quantification SpdyE vs GAPDH
2.5
2
1.5
1
0.5
0
231 Cntl
231 Spy1 KO
C
Relative Quantification CycA vs GAPDH
1.6
1.4
1.2
1
0.8
0.6
0.4
0.2
0
231 Cntl
231 Spy1 KO
D
Relative Quantification CycB vs GAPDH
2
1.5
1
0.5
0
231 Cntl
231 Spy1 KO
E
Relative Quantification CycD vs GAPDH
2
1.5
1
0.5
0
-0.5
231 Cntl
231 Spy1 KO
F
Relative Quantification CycE vs GAPDH
2
1.5
1
0.5
0
-0.5
231 Cntl
231 Spy1 KO

Figure S2

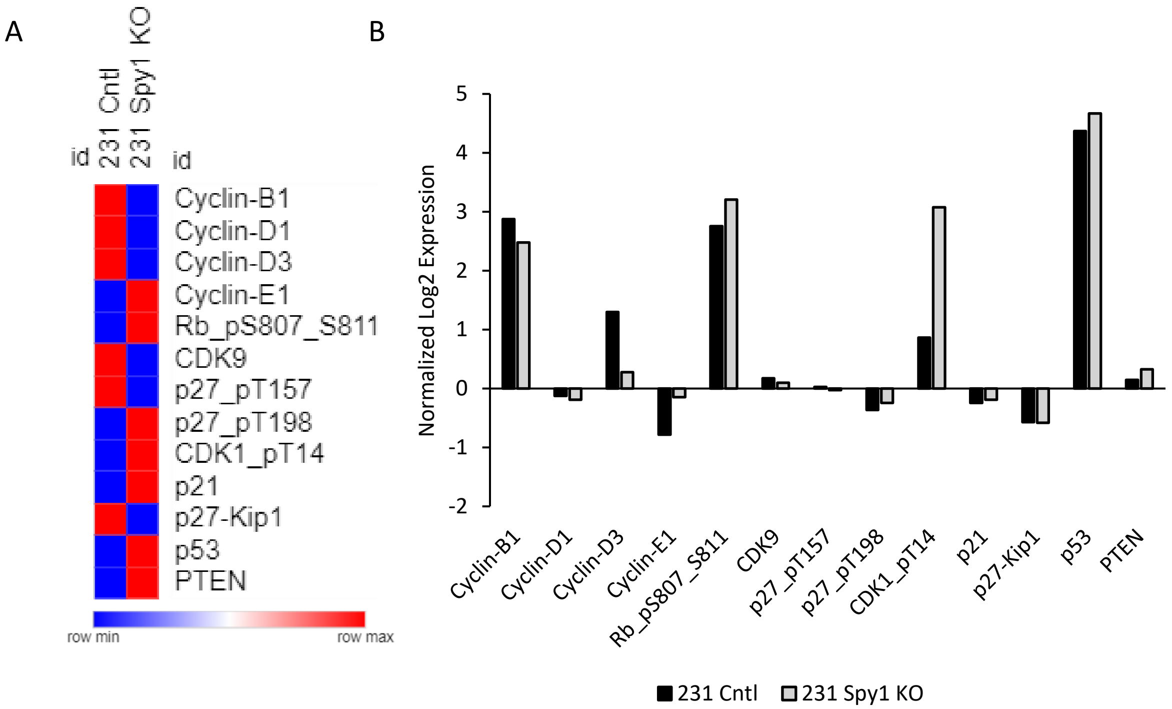


C

| | Cyclin-B1 | Cyclin-D1 | Cyclin-D3 | Cyclin-E1 | Rb_pS807_S811 | CDK9 | p27_pT157 | p27_pT198 | CDK1_pT14 | p21 | p27-Kip1 | p53 | PTEN |
|---|---|---|---|---|---|---|---|---|---|---|---|---|---|
| **231 Cntl** | 2.873117 | -0.12538 | 1.301139 | -0.7793 | 2.758009 | 0.17259 | 0.030769 | -0.36347 | 0.86348 | -0.24143 | -0.5697 | 4.368249 | 0.145373 |
| **231 Spy1 KO** | 2.477431 | -0.18959 | 0.276733 | -0.14407 | 3.207619 | 0.09644 | -0.02672 | -0.24601 | 3.076604 | -0.19113 | -0.58055 | 4.668731 | 0.328584 |

Average expression positive regulators - Average expression negative regulators = Proliferation Score

Figure S3

A

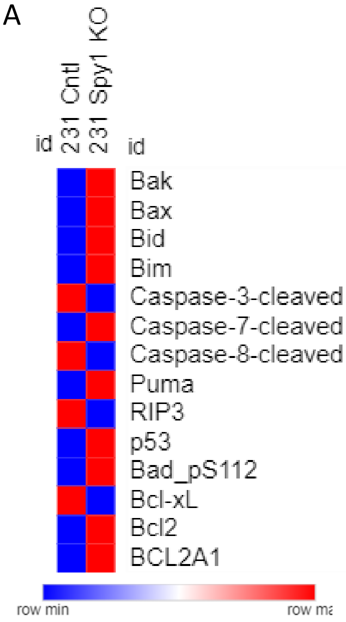


B

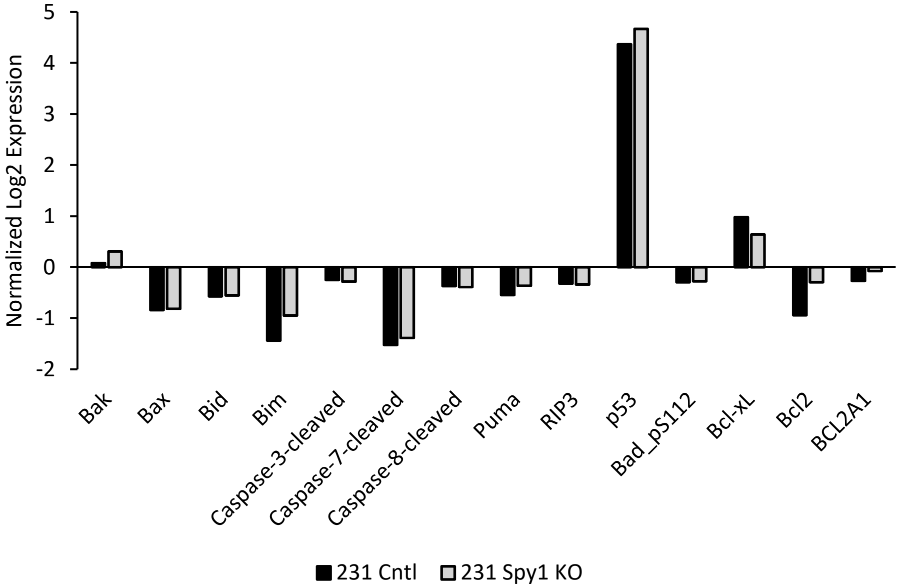


C

| | Bak | Bax | Bid | Bim | Caspase-3-cleaved | Caspase-7-cleaved | Caspase-8-cleaved | Puma | RIP3 | p53 | Bad_pS112 | Bcl-xL | Bcl2 | BCL2A1 |
|---|---|---|---|---|---|---|---|---|---|---|---|---|---|---|
| 231 Cntl | 0.080167 | -0.84084 | -0.57423 | -1.43865 | -0.25004 | -1.52524 | -0.3689 | -0.54457 | -0.32069 | 4.368249 | -0.29654 | 0.979435 | -0.93912 | -0.26713 |
| 231 Spy1 KO | 0.310104 | -0.81308 | -0.55286 | -0.94936 | -0.28069 | -1.38756 | -0.3917 | -0.36214 | -0.34026 | 4.668731 | -0.2745 | 0.637876 | -0.2938 | -0.07775 |

Average expression positive regulators - Average expression negative regulators = Apoptotic Score

Figure S4

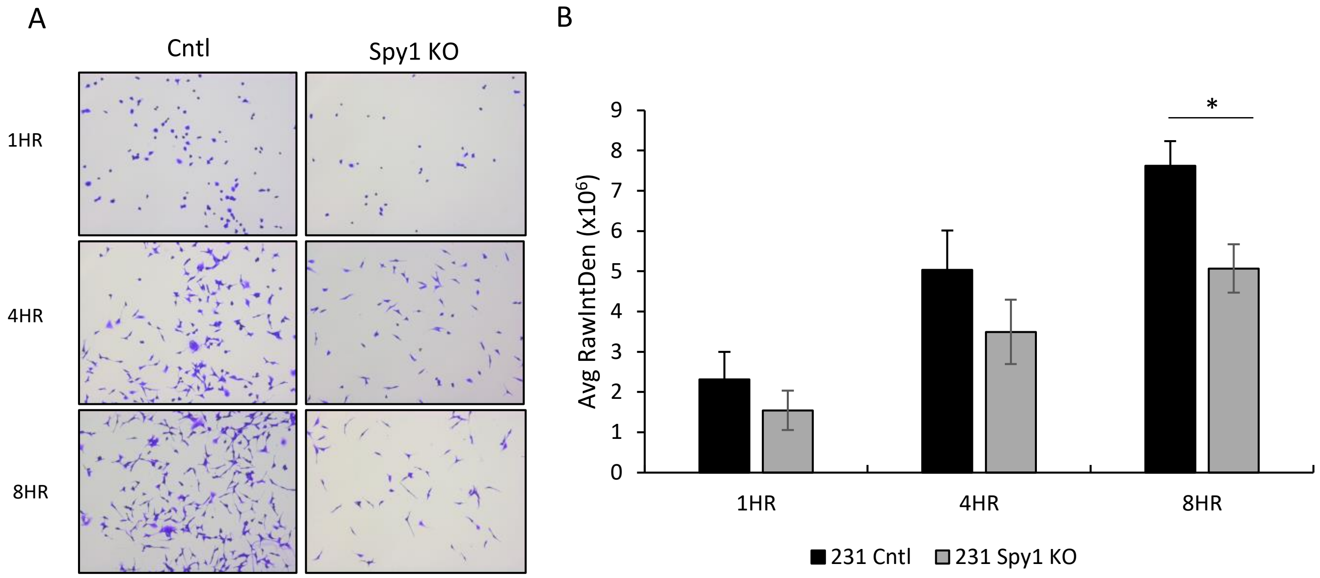
A
Cntl
Spy1 KO
1HR
4HR
8HR
B
Avg RawIntDen (x10^6)
0
1
2
3
4
5
6
7
8
9
*
1HR
4HR
8HR
231 Cntl
231 Spy1 KO

Figure S5

A

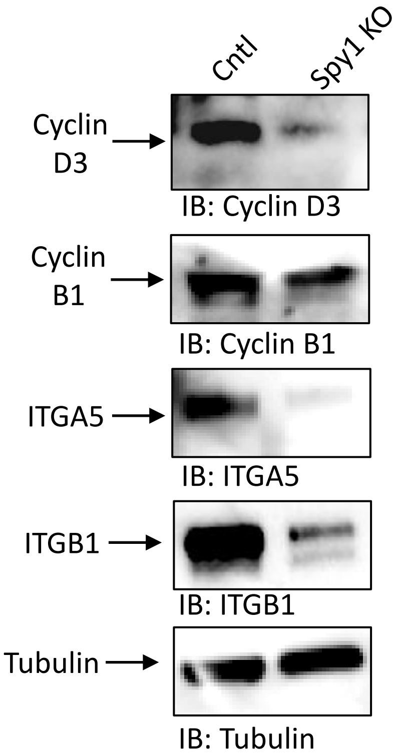
Cntl
Spy1 KO
Cyclin D3
IB: Cyclin D3
Cyclin B1
IB: Cyclin B1
ITGA5
IB: ITGA5
ITGB1
IB: ITGB1
Tubulin
IB: Tubulin

B

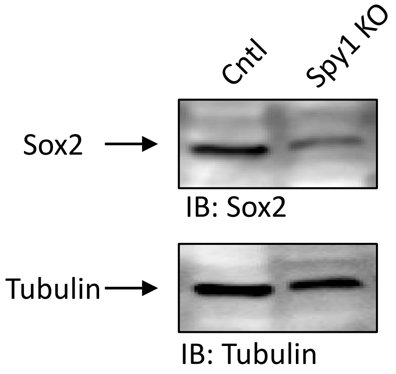
Cntl
Spy1 KO
Sox2
IB: Sox2
Tubulin
IB: Tubulin

Figure S6

A

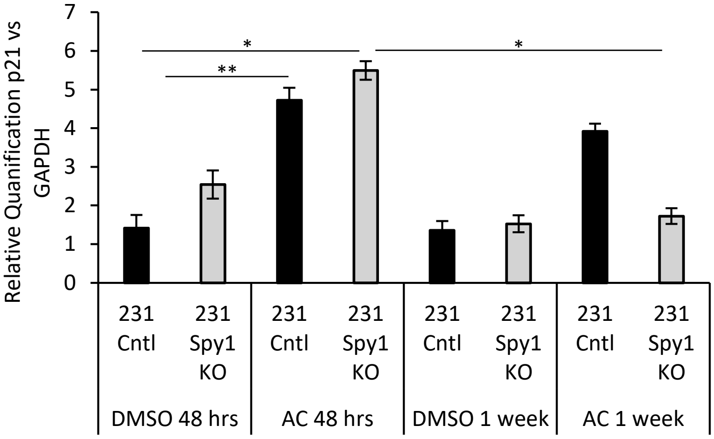

Relative Quanification p21 vs GAPDH
7
6
5
4
3
2
1
0
*
**
*
231 Cntl
231 Spy1 KO
231 Cntl
231 Spy1 KO
231 Cntl
231 Spy1 KO
231 Cntl
231 Spy1 KO
DMSO 48 hrs
AC 48 hrs
DMSO 1 week
AC 1 week

### Supplemental Figure Legends

**Figure S1: Cyclins and Speedy/RINGO family members do not compensate for loss of Spy1 expression.** qRT-PCR analysis examining levels of **A)** SpdyC, **B)** SpdyE, **C)** Cyclin A, **D)** Cyclin B, **E)** Cyclin D1 and **F)** Cyclin E corrected for total GAPDH levels. n=3; Error bars represent SE; Student's T test.

**Figure S2: Proliferation score of Spy1 knockout cells. A)** Heat map of log2 expression profiles of cell cycle regulators from RPPA analysis. **B)** Graphical depiction of normalized log2 expression profiles of cell cycle regulators from RPPA analysis. **C)** Log2 expression of cell cycle regulators used in calculation of proliferation score where green represents positive drivers of proliferation and red represents negative drivers of proliferation.

**Figure S3: Apoptotic score of Spy1 knockout cells. A)** Heat map of log2 expression profiles of apoptotic regulators from RPPA analysis. **B)** Graphical depiction of normalized log2 expression profiles of apoptotic regulators from RPPA analysis. **C)** Log2 expression of apoptotic regulators used in calculation of apoptotic score where green represents positive drivers of apoptosis and red represents negative drivers of apoptosis.

**Figure S4: Spy1 knockout reduces adhesion. A)** Representative images of MDA-MB-231 adhesion assay of Spy1 knockout and control cells on tissue culture plates at 1hr, 4hr and 8hr timepoints. Images were taken at 10X magnification. Scale bar represents 200μm. **B)** Graph represents quantification of MDA-MB-231 Spy1 knockout cells compared to MDA-MB-231 control cells adhered to the plate at respective timepoints. n=3. Error bars represent SE; Student's T test. $*p < 0.05$, $**p < 0.01$, $***p < 0.001$

**Figure S5: Loss of Spy1 results in decreased expression of prognostic markers *in vivo*. A)** Levels of cyclin D3, cyclin B1, integrin β1, integrin α5 and **B)** Sox2 were assessed via western blot analysis in MDA-MB-231 control and Spy1 knockout tumours collected at end point from NOD/SCID mice. Tubulin was used as a loading control. n=2.

**Figure S6: Decreased expression of p21 following treatment with Spy1 knockout. A)** qRT-PCR analysis of p21 levels corrected for GAPDH of MDA-MB-231 control and Spy1 knockout cells treated with AC. A=doxorubicin, C=cyclophosphamide; n=3. Error bars represent SE; Student's T test. $*p < 0.05$, $**p < 0.01$, $***p < 0.001$
